# Extending the IICR to complex non-stationary structured models

**DOI:** 10.1101/2024.10.21.619462

**Authors:** Alexane Jouniaux, Armando Arredondo, Simon Boitard, Lounès Chikhi, Olivier Mazet

**Affiliations:** Institut des Mathématiques de Toulouse, Université de Toulouse, Institut National des Sciences Appliquées, 31077 Toulouse, France; CBGP - Centre de Biologie et Génétique des Populations, Université de Montpellier, CIRAD, INRAE, Institut Agro, IRD, Montpellier, France; Centre de Recherche sur la Biodiversité et l’Environnement, Université de Toulouse, CNRS, 31062 Toulouse, France; CE3C - Centre for Ecology, Evolution and Environmental Changes, Departamento de Biologia Animal, Faculdade de Ciências, Universidade de Lisboa, 1749-016 Lisboa, Portugal; Instituto Gulbenkian de Ciência, Oeiras, Portugal

**Keywords:** demographic inference, coalescent theory, IICR, structured coalescent, non-stationary models, glue matrices

## Abstract

Population genetic studies use genetic data to understand aspects of the evolutionary history of species, particularly to infer their demography. Here, we extend the non-stationary structured coalescent framework to complex models where the parameters of the model, and thus the state space, change between stationary periods. To do this, we introduce glue matrices that allow us to map one state space to another, hence enabling the computation of the Inverse Instantaneous Coalescence Rate (IICR) using Q-matrices of different dimensions. This approach allows us to study models where the number of demes changes due to extinction or foundation. Our analysis confirms that interpreting IICR curves as indicators of population size changes can be misleading. However, we also show that there are cases where new deme foundations can generate an increase in the IICR (forward in time). We explore the impact of changes in migration rates, timing of deme creation or deletion, and sampling strategies on the IICR, identifying several counter-intuitive results. For instance, we find that the IICR starts to decrease several generations before any change in the number of demes, as if the population “knew” something was about to happen. It is particularly unexpected when a deme is founded and the population is only increasing. Our results underscore the need for careful interpretation of PSMC and similar curves, which are often used and interpreted as population size changes. By presenting examples of IICRs for various transitions and sampling scenarios, we emphasize the importance of nuanced approaches in population genetic studies.

## Introduction

The reconstruction of the demographic history of populations and species using genetic data remains a complex yet essential challenge for evolutionary and conservation biologists, as well as for theoreticians, providing insights into the evolutionary history of species (Scerri *et al*. 2018; Liu and Fu 2015; Johri *et al*. 2021; Hey and Machado 2003; Goldstein and Chikhi 2002). The dynamic interplay between demographic shifts (expansions and contractions) and environmental changes (habitat fragmentation, extension, and contraction) shapes not only historical population sizes and connectivity patterns (Wasserman *et al*. 2012; Gubili *et al*. 2017; Selwood *et al*. 2015; Young *et al*. 2018) but also their spatial distribution through time (Haddad *et al*. 2015; Cushman 2006; Pacifici *et al*. 2020; Wolf and Ripple 2017; Finn *et al*. 2023) and current levels of genetic diversity (Alcala *et al*. 2019; Dures *et al*. 2019; Cabe 1998; Nei *et al*. 1975). Many inference methods assume a single panmictic population, viewing demographic history only as a series of population size changes (Sheehan *et al*. 2013; Beaumont 1999; Chevalet and Nikolic 2010; Bunnefeld *et al*. 2015). However, it has become clear that multiple factors, such as sampling strategies and population structure (Mazet *et al*. 2016; Nielsen and Beaumont 2009; Chikhi *et al*. 2010; Beaumont 2004; Städler *et al*. 2009; Wakeley 2001, 1999), must be taken into account for accurate inference. To address these complexities, various methods have been developed to elucidate both recent and ancient population structures (Al-Asadi *et al*. 2019; Gutenkunst *et al*. 2009; Harris and Nielsen 2013; Palamara and Pe’er 2013; Link *et al*. 2017; Beeravolu *et al*. 2017; Raj *et al*. 2014; Pritchard *et al*. 2000; François and Jay 2020; Joseph and Pe’er 2019; Arredondo *et al*. 2021). These approaches have been applied not only to diverse endangered species (Teixeira *et al*. 2021) but also to human populations. In this study, we focus on methods based on the structured coalescent, particularly within the framework of the Inverse Instantaneous Coalescence Rate (IICR), as introduced by Mazet *et al*. (2016).

In a few words, the IICR is a time- and sample-dependent function of the distribution of coalescence times for the demographic model of interest. It provides the inverse rate of coalescence events for a sample of size *k* haploid genomes. It was originally defined for *k* = 2 (Mazet *et al*. 2016) but has now been extended (Chikhi *et al*. 2024) to any *k* ≥ 2 (for diploid organisms we expect *k* to be even and to correspond to *p* diploid individuals, where *k* = 2*p*, but the theory is valid for any *k*). In a panmictic setting, the IICR corresponds to the coalescent effective size *N*_*e*_ but its interpretation is much less clear under structured models. For *k* = 2 the IICR can be estimated using the PSMC method of Li and Durbin (2011) or the MSMC method of Schiffels and Durbin (2014). Theoretical work on the IICR suggests that under structured models the PSMC and MSMC curves may exhibit humps and decreases even when population sizes did not change, and such humps could be misinterpreted as signals of population size changes. Given that genomic studies focusing on endangered species rarely achieve large sample sizes, methods that require the genome of only one or a few individuals such as the PSMC and MSMC methods, have been very popular since they were developed in the last decade and a half. It is thus important to comprehend the properties of the IICR and what it can tell us about the demographic history of populations, and what it cannot.

In a previous study, Rodríguez *et al*. (2018) employed Markov chain theory to derive the IICR for structured step-wise stationary models, where the number of demes was kept constant and migration changed in a step-wise manner (*i*.*e*. migration was assumed to be constant for certain time periods, but was allowed to change between periods). By constructing a Q-matrix for each stationary model, they showed that the IICR could be made computationally accessible, overcoming challenges posed by the absence of analytical formulas for the density of coalescence times in complex structured models. The size of the different matrices is determined by the number of states, which is the number of configurations of the sampled genes at any time, which itself depends on the sample size and the structured model. This framework thus allowed to study both stationary and non-stationary models, provided the number of states in each model was the same, as the computation of the IICR required the multiplication of the Q-matrices.

This theory has been successfully used to develop an inferential method called SNIF (Structured Non-stationary Inferential Framework Arredondo *et al*. (2021)) which allows to infer the parameters of step-wise stationary *n*-island models where *n* is inferred but does not change between stationary periods. In other words, while the framework allowed for changes in connectivity, it did not permit a dynamic change in the number of states, such as those implied by a change in the number of demes. Rodríguez *et al*. (2018) showed that it was possible to change the number of demes using an *ad-hoc* solution where all matrices are expanded to the size of the largest matrix required by one of the stationary periods, but they did not provide a general solution allowing to change the number of demes dynamically.

In response to this limitation, our study introduces glue matrices which, in the context of Markov chains, allow to map the states of the coalescent process before and after a change in the number of states. In this article, we also propose a model for this state mapping and demonstrate how to compute a glue matrix accordingly. We then examine some properties of the non-stationary IICRs it produces. Finally, we explore processes such as colonization, splits, habitat fragmentation, and habitat loss, thereby improving our understanding of population history modeling over time.

## The structured coalescent and the IICR

The distribution of coalescence times is significantly shaped by population structure. Over the past few decades, numerous theoretical studies have improved our understanding of this distribution within the framework of population structure modelling. In this section, we specifically present the structured coalescent model developed by Herbots (1994) and Notohara (1990). This model uses Markov chain theory to compute both the probability density function (pdf) and cumulative distribution function (cdf) of coalescence times as a function of the demographic model and sampling scheme, and allows the presentation of the Inverse Instantaneous Coalescence Rate (IICR) as originally outlined by Mazet *et al*. (2016) for two lineages.

### Structured Coalescent

We model populations that deviate from panmixia using the structured coalescent (Herbots 1994; Notohara 1990). We assume that a metapopulation is subdivided into a finite number *n* of demes (or islands), each of which is assumed to evolve under a haploid Wright-Fisher model. The islands can be connected and exchange genes between them in an arbitrary manner, even though most studies have focused on symmetrical models such as the *n*-island or stepping-stone models. At each generation, the model involves two key steps: reproduction and migration, ensuring the population size of each deme remains constant. During the reproduction step, the entire generation is replaced by the subsequent one, with each gene randomly selecting its parent from the preceding generation.

We define the number of haploid genes in deme *i* as *N*_*i*_ = 2*s*_*i*_ *N* where *s*_*i*_ is the relative deme size (*i*.*e*. 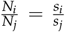 for every other deme *j*) and *N* is assumed to be large. During the migration step forward in time, each gene has a probability *q*_*ij*_ to migrate from deme *i* to deme *j*. Thus, the number of genes migrating from *i* to *j* is *q*_*ij*_*s*_*i*_ and those migrating from *j* to *i* is *q*_*ji*_*s*_*j*_. Notably, these probabilities are assumed to be small, on the order of 1/*N*. Two different strategies have been developed to maintain constant population sizes in each deme. Herbots (1994) assumes conservative migration where

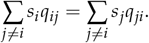

This means that the number of genes leaving island *i* is equal to the number of genes arriving in island *i*, allowing to maintain the size of island *i* constant.

In contrast, Notohara (1990) suggested performing the migration step before reproduction, allowing for non-conservative migration and restoring each deme’s population size to its original value. Under this model, the islands do not evolve under a Wright-Fisher model anymore, but a recent study by Kozakai *et al*. (2016) demonstrated that both strategies lead to the same limit process.

We then introduce the backward migration rate 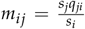 as the proportion of lineages or genes from island *i* that originated from island *j* in the previous generation. We assume that ∀*i, j* ∈ 1, …, *n* with *i*≠ *j*, the sequence (2*Nm*_*ij*_)_*N*_ is an increasing sequence such that

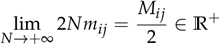

and denote *M*_*i*_ = ∑_j≠*i*_ *M*_*ij*_. A convenient way to represent all migration parameters *M*_*ij*_ is to put them in a migration matrix *M* ∈ M_*n*×*n*_, with *M*_*ij*_ being the coefficient at line *i* and column *j*.

We can then sample *k* haploid genes from the present population and trace back the genealogy of their genes until we identify a common ancestor between any pair of lineages in the sample. The set of all possible states (or configurations) of *k* lineages is denoted *E*_*k,n*_ and defined as 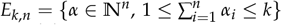. Now, we can define the vector *α*_*N*_ (*τ*) ∈ *E*_*k,n*_ as the configuration of the distinct ancestors of the sample of *k* genes *τ* generations ago. Here, each component *α*_*i*_ represents the number of lineages in deme *i* for each 1 ≤ *i* ≤ *n*. Scaling time in 2*N* generations, we obtain the ancestral process (*α*_*N*_ (⌊2*Nt*⌋))_*t*≥0_ which converges to a Markov process known as the structured coalescent (Herbots 1994), characterized by the matrix *Q*:

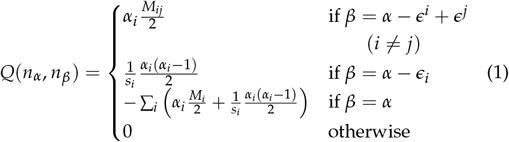

where *α, β* ∈ *E*_*k,n*_ and *ϵ*^*i*^ is the vector whose components are all 0 except for the *i*-th one that is 1. We define *nα* and *n*_*β*_ as the indices of the row of configuration *α* and *β* respectively. The first case in (1) corresponds to a migration event of one of the *α*_*i*_ lineage from deme *i* to deme *j*, the rate of each migration being *M*_*ij*_/2. The second case stands for a coalescence event, it can occur only if *α*_*i*_ ≥ 2. The binomial coefficient *α*_*i*_ (*α*_*i*_ −1)/2 represents the number of possible pairs among the *α*_*i*_ lineages and 1/*s*_*i*_ is the coalescence rate for each pair of lineages in the island *i*.

Sometimes, the symmetry of the model allows for the simplification of the Q-matrix. For example, in the *n*-island model, if we consider a sample of size *k* = 2, all states can be summarized into three distinct states: the two lineages are in the same island (denoted as *s*), in different islands (denoted as *d*), or they have coalesced. We then obtain the following Q-matrix:

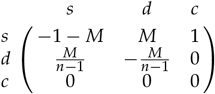

However, in this work, we will not consider such symmetry, and we will always explicitly write the entire Q-matrix using Equation 1.

The structured coalescent described above is general on that it can be used to study any model of population structure and compute the corresponding IICRs.

### Inverse Instantaneous Coalescence Rate (IICR)

The IICR (Mazet *et al*. 2016) was first defined by considering the coalescence times of two lineages. It can be derived analytically in the case of the *n*-island model and otherwise be computed for any structured model using the theory of Markov chains described previously. It has been extended to *k* = 3 (Grusea *et al*. 2019), and then to *k* genes (Chikhi *et al*. 2024). In this section, we describe the latter case.

We sample *k* genes at the present (*t* = 0) and trace their ancestral lineages back in time until the first coalescence event among them. These lineages are in configuration *α*. By employing the theory of Markov chains within the framework of the structured coalescent, we have the necessary tools to compute the probability distribution of the coalescence time, denoted as *T*_*k,α*_.

We derive the transition probabilities between two states by exponentiating the transition matrix *Q* to obtain the transition semigroup of the corresponding Markov process, expressed as *P*_*t*_ = *e*^*tQ*^ (for a comprehensive introduction to Markov chains, see for instance Norris (1998)). This provides the transition probabilities, representing the likelihood that *k* lineages in configuration *α* at time *t* = 0 will be in state *β* at time *t*. Each coefficient corresponds to

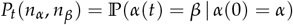

In particular, we are interested in the probability that at least one pair among the *k* lineages sampled in configuration *α* at the present time has already coalesced by time *t*. This probability is denoted as *P*_*t*_ (*n*_*α*_, *n*_*c*_), where *n*_*c*_ is the column of *P*_*t*_ representing the coalescence state *c* and *n*_*α*_ the column of *P*_*t*_ representing the configuration *α*. Several coalescence states exist, each one corresponding to an element of the set 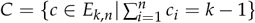.

Therefore, if we define *T*_*k,α*_ as the coalescence time for *k* lineages sampled in configuration *α*, we can compute the cdf of this random variable using the transition semigroup as follows:

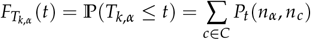

The pdf of *T*_*k,α*_, denoted by 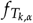, is defined as the derivative of 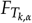. This can be computed easily with *P*_*t*_ using the property 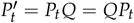 (Kolmogorov forward equation), where *P*^′^ is a matrix whose elements contain the derivative of the corresponding elements in *P*_*t*_. In conclusion, we have the following:

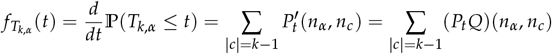

The instantaneous coalescence rate at time *t* is defined as the probability that two lineages that have not yet coalesced at time *t* (denominator) will do so in an infinitesimal interval of time starting at *t* (numerator): 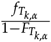 (Mazet *et al*. 2016; Chikhi *et al*. 2024).

We take the inverse of this ratio to obtain the IICR, which corresponds to the effective size of the population under the panmixia assumption. The *IICR*_*k,α*_ can be expressed in terms of *P*_*t*_ and the transition rate matrix using the following equation:

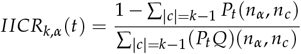

## Non-Stationary Structured Coalescent (NSSC)

The previous descriptions focused on stationary models, where the population structure remains constant over time. Here, we introduce a new level of complexity by allowing non-stationarity in the considered models. When population structure undergoes alterations, it may lead to a change in the number of states in the coalescent process. Such changes can introduce asymmetry in migration rates or deme sizes, or a variation in the number of demes. We thus need to consider each island and its associated states independently. As mentioned in section The structured coalescent and the IICR, we will not simplify the Q-matrix.

Initial efforts by Rodríguez *et al*. (2018) to account for non-stationarity assumed that the number of states remains constant. We will begin by presenting the mathematical formulation of the non-stationary structured coalescent model, assuming a constant number of states. Subsequently, we will explain how to extend this formulation to non-stationary models involving a change in the number of states.

We assume that the population has a certain structure at the present time (*t* = 0), described by the parameter vector (*n*_0_, *s*_0_, *M*_0_), where *n*_0_ is the number of islands, *s*_0_ is the vector containing the sizes of the demes, and *M*_0_ is the migration matrix containing the rates of migration between these sub-populations. As we go back in time, the population undergoes configuration change at time *T*. This altered configuration is described by the parameter vector (*n*_1_, *s*_1_, *M*_1_). We denote *Q*_0_ as the transition rate matrix of the Markov chain for 0 ≤ *t* ≤ *T*, and *Q*_1_ as the transition rate matrix for *t* > *T*.

In the different examples we will present in this work, migration rates between islands are symmetrical and maintained constant in the sense that the existing migration rates remain the same, and the added ones are equal to the existing ones. However, it is possible to model structures that transition from symmetric to asymmetric migration rates. We do not impose any constraints on these migration rates. It is important to note that adding a deme introduces additional complexity to the model at the level of migration rates. Specifically, if we add a new neighbor to an existing island, the existing island will send more genes at each migration step than before. To ensure that the “total” migration of an island remains constant, we need to adjust our migration rates accordingly when defining our models.

We can take for example the transition between a 2-island and a 3-island model and set *M* = *M*_*i*_, meaning that each island will receive *M* genes at each migration step (*M* = 2 for the 3-island and *M* = 1 for the 2-island). We clearly see that when a deme is founded or go extinct, the number of immigrants arriving in the other islands increases or decreases. If we wanted to keep it constant, we would have to set the migration rates *M*_*ij*_ to 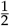 in the 3-island model (or to 2 in the 2-island).

### NSSC with constant number of states

We consider the non-stationary model associated with the parameter vectors (*n*_0_, *s*_0_, *M*_0_) and (*n*_1_, *s*_1_, *M*_1_), which correspond to a change in population configuration that does not involve a modification in the number of states (*n*_0_ = *n*_1_). This implies that *Q*_0_ and *Q*_1_ have the same size.

The semigroup of a Markov process satisfies the following identity, called the semigroup property:

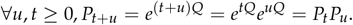

This property is used to extend the structured coalescent model to non-stationary models.

We first assume that the number of states of the coalescent process is constant over time. We can then define the matrix 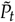 as the transition semigroup of the Markov chain for this non-stationary process:

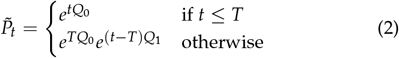

Again, we can compute the cdf of *T*_*k,α*_:

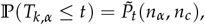

as well as the pdf of *T*_*k,α*_:

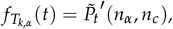

where:

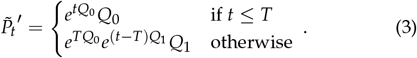

We can extend these results to any number of parameter changes.

### NSSC with changing number of states

Several phenomena can lead to a change in the number of states, such as introducing asymmetry in migration rates or deme sizes, or creating or removing a deme. By considering each island and its associated states independently and writing the entire Q-matrix explicitly, we already account for potential asymmetries in migration rates and deme sizes. However, when the number of demes changes, it requires a precise definition of how we map the new states with the previously existing ones. This requires accurate mathematical modeling to describe these operations.

First, we describe in a forward perspective what happens when the number of islands changes. We start with a certain number of demes in the remote past. At some point, some of them may go extinct, meaning their genes cannot be in the present sample, neither contribute to its genealogy. Alternatively, new islands may be founded and connected to the existing ones. We will refer to the first one as the “deme extinction” scenario and the second one as the “deme foundation” scenario.

But since we are in the coalescent framework, we frequently adopt a backward perspective, even though it may not be the most intuitive. To maintain clarity about the chosen perspective in the following, we will use the terms “deme suppression” and “deme addition” to specifically refer to scenarios from the backward perspective. The correspondence between forward and backward perspectives is as follows:

|  |
| --- |
| Forward $\iff$ Backward |
| Deme foundation $\iff$ Deme suppression |
| Deme extinction $\iff$ Deme addition |

The scenario of a deme extinction corresponds to having fewer demes in the present and recent past than in the remote past. In this case, all states are called “persistent”, meaning that the states continue to exist after the change in configuration from a backward perspective, despite the changes in the number of states.

Conversely, in the case of a deme foundation, we have more demes in the present and recent past than in the remote past. Some of the states are no longer persistent because they were associated with an island that is no longer part of the structure. This implies that we must ensure the ancestral lineage of our sample continues to exist after the change in the past. Therefore, we assume that genes present in the removed deme move to a neighboring island. Each lineage in the deleted deme has a probability, depending on the migration rate, to move to a specific neighboring island.

A crucial assumption in our modeling of population suppression is that lineages in a suppressed deme must move to a neighboring island that continues to exist in the past. This ensures that we can always define a coherent mapping between states, without the complexity of allowing genes to move to a more distant island, which may not be biologically relevant. Extra caution must be used, especially in the case of multiple deme suppressions, to ensure that we are not deleting an island and its only neighbors. An example where this condition is not met is shown in Figure 1, where islands 3 and 4 are suppressed and then island 4 does not have an existing neighbour in the present.

**Figure 1.**
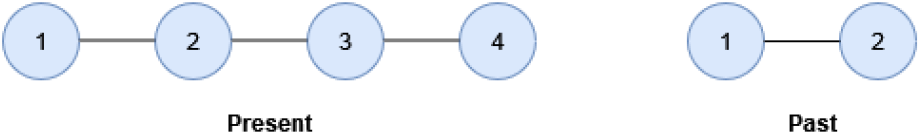
Example of an impossible transition in which we go from a linear stepping-stone with two demes to a linear stepping-stone with four demes. The deme 4 does not have existing neighbour in the present.

Mathematically, if we change the number of demes from *n*_0_ to *n*_1_ looking backward in time and consider a sample of size *k* = 2, the Q-matrices *Q*_0_ and *Q*_1_ have different sizes 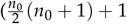 and 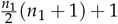 respectively, where 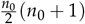 is the number of possible transitions between states, equals to the number of pairs of islands, and the +1 stands for the coalescence state). It is then impossible to compute the product 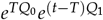 in Equation 2 and 3.

The solution we propose is to match the states of the coalescent processes associated with the parameter-vectors (*n*_0_, *s*_0_, *M*_0_) and (*n*_1_, *s*_1_, *M*_1_) via a so-called glue matrix, *G*, of size 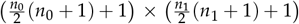 that will allow the multiplication 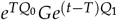.

In the following, we will only consider the case of a sample of size 2, therefore *T*_2,*α*_. The set of all possible states will be thus 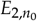 until the configuration change and 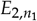 after.

#### Definition of the glue matrix

Thereafter, the notation (*ij*) will refer to the state of two lineages and means that one lineage is in island *i* and the other in island *j*. It corresponds to a configuration *α* ∈ *E*_2,*n*_ in which the components *i* and *j* are set to 1, and others are set to 0. Because we assume that the lineages are indistinguishable, so are the states (*ij*) and (*ji*); therefore, in the following, we will always suppose that *i* ≤ *j*. The glue matrix *G* will explicitly provide the “instantaneous” probability of changing from one state in 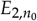 to another state in 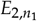 at the time of the transition *T*. These instantaneous probabilities must be described accurately because certain states will be removed while others will be added due to the addition or suppression of demes.

More specifically, suppose that we have two states, (*ij*) and (*kl*) with (*ij*) being a possible configuration in the set 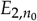 and (*kl*) a possible configuration in the set 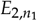. We denote by *n*_(*ij*)_ the index of the row of configuration (*ij*) and *n*_(*kl*)_ the index of the column of configuration (*kl*) in the glue matrix. Then, we have *G*(*n*_(*ij*)_, *n*_(*kl*)_) = ℙ *α*(*T*) = (*kl*) | *α*(*T*^−^) = (*ij*). The probabilistic meaning of the glue matrix is explained more extensively in the Supplementary Material. We denote by (*i*•) the configuration where one lineage is in deme *i* and the other is in any deme. For the sake of clarity, we will use the notation ℙ(*i* → *k*) for the probability that a lineage in island *i* moves to island *k* (*i*.*e*. ℙ [*α*(*T*) = (*k*•) | *α*(*T*^−^) = (*i*•)]).

#### Computation of the instantaneous probabilities

We can now describe how we compute the instantaneous probabilities according to our modeling (see beginning of section The structured coalescent and the IICR). We want to stress that different hypotheses of modeling will induce different computations. As defined previously, *M*_*ij*_ is the **backward** migration rate from *i* to *j*. Migration between islands *i* and *j* can be asymmetric, *M*_*ij*_≠ *M*_*ji*_. We say that island *j* is a neighbour of island *i* at time *t* if *M*_*ij*_ > 0. We can then write *j* ∈ *N*_*t*_ (*i*) where *N*_*t*_ (*i*) is the set of neighbours of island *i* at time *t* and we denote by |*N*_*t*_ (*i*)| the number of neighbours of the island *i* at time *t*.

One can customize the glue matrix according to their own modelling hypotheses, with the only constraint being that the matrix must be stochastic, meaning that all its entries are non-negative and each row sums to one. The results we will present in the next section are entirely dependent on the choice of glue matrix. In the Supplementary Material, we present another modeling approach for which we could not find a biological explanation but produces interesting results.

We start by addressing the coalescence state that will simply map to itself. In other words, we have *G*(*n*_*c*_, *n*_*c*_) = 1. Another straightforward case is that of a persistent state, where lineages do not move. It will also map to itself, denoted as *G*(*n*_(*ij*)_, *n*_(*ij*)_) = 1.

When a state does not persist after the change, one or both lineages must move to a neighboring island. The probability that a lineage in island *i* moves to island *k* is computed as follows:

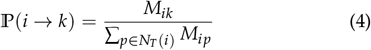

where *N*_*T*_ (*i*) is the set of neighboring islands that persist after the change. In this equation, the numerator represents the migration rate to island *k*, and the denominator represents the total migration rate from island *i* to neighboring islands.

If only one lineage needs to move from island *i* to a neighbour island *k*, which corresponds to a transition from state (*ij*) to (*kj*), we apply the formula (4).

However, when both lineages must move to effect the transition from (*ij*) to (*kl*), two possibilities arise:

- If we are in the case where *i* = *j*, that corresponds to a change from state (*ii*) to (*kl*) with *k*≠ *l*, we have: *G*(*n*_(*ii*)_, *n*_(*kl*)_) = 2 × ℙ(*i* → *k*) × ℙ(*i* → *l*), where we have a factor 2 because each lineage moves independently.
- Otherwise, if we change from state (*ij*) to state (*kl*) with *k* ∈ *N*_*t*≥*T*_ (*i*) and *l* ∈ *N*_*t*≥*T*_ (*j*), we have: *G*(*n*_(*ij*)_, *n*_(*kl*)_) = ℙ(*i* → *k*) × ℙ(*j* → *l*).

##### Reference example

In this study, we use the reference example of Figure 2 to illustrate how glue matrices are constructed and to study the properties of the corresponding non-stationary IICR. Specifically, we examine transitions between a two-island and a three-island model with the migration rates between all the different islands set to 1, which means that *M*_*ij*_ = 1 ∀*i*≠ *j*.

**Figure 2.**
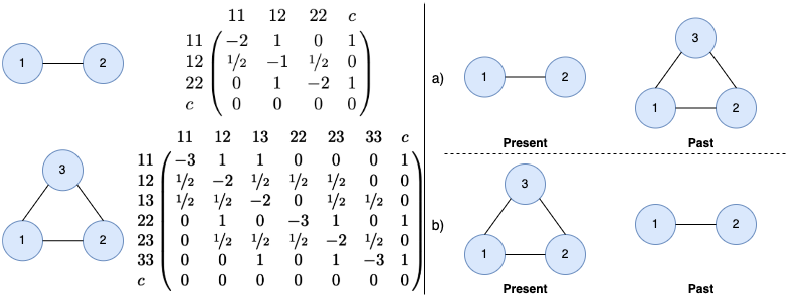
Reference example with 2- and 3-island models, showing their Q-matrices (left) and the transitions possible between them (right). (a) corresponds to a deme extinction/deme addition and will be referred to as Example 1, and (b) corresponds to a deme foundation/deme suppression and will be referred to as Example 2.

##### Example 1: Deme addition backward in time

We study first the backward transition from a 2-island in the present to a 3-island model in the past (see Figure 2 a)). Because all states of this non-stationary process are persistent, the glue matrix is straight-forward to compute. We simply have that *G*(*n*_(*ij*)_, *n*_(*kl*)_) = 1, where (*ij*) equals to (*kl*). It results in the following glue-matrix:

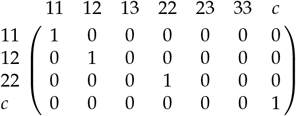

##### Example 2: Deme suppression backward in time

We look now at the backward transition from 3-island in the present to 2-island model in the past (see Figure 2 b)), which is more complex. All the transitions between persistent states ((11),(12),(22)) are set to 1.

If the two lineages are in the state (13), the lineage in island 3 can move to island 1 or 2 with the same probability because of the symmetry of migration rates. Using the formula 4, we have 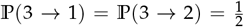. The lineage in island 1 remains where it is, that is ℙ(1 → 1) = 1. It results that 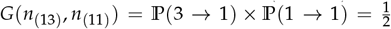 and 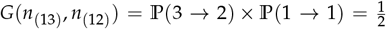. The case where the two lineages are in the state (23) is analogous.

If the two lineages are in the state (33), both lineages must move to either island 1 or island 2. We use again the formula 4 that gives 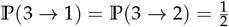. We will obtain then

- 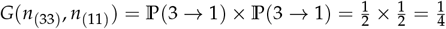
- 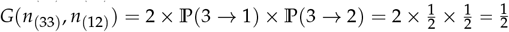
- 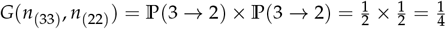

We obtain the following glue-matrix (still with the example of Figure 2):

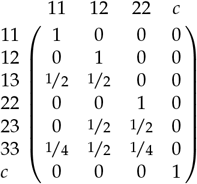

## Results

We have now developed a general framework to compute IICRs for a wide range of non-stationary models, where the number of states varies between stationary periods. In this study, we focus on models that involve changes in the number of demes, hence allowing us to study processes such as colonization, extinction and fragmentation. We also investigate how various parameters of the model influence the IICR.

### IICR of the reference example

Figure 3 illustrates the IICR for the reference example detailed in section Reference example, depicting the two possible transitions (deme addition and suppression) and their corresponding real population size evolution. Additionally, the IICRs of the stationary 2-island and 3-island models are included for comparative analysis. The two lineages in our sample originate from the same island (island 1), as evidenced by the characteristic S-shaped curves of the IICR for samples taken from one deme of an *n*-island models. (Refer to Supplementary Figure 9 for sampling across different islands.) As previously demonstrated by Mazet *et al*. (2016), it is noteworthy that the IICRs bear little resemblance to actual population sizes (dotted lines), highlighting the limitations of using IICRs as a direct proxy for population size evolution under simple structured models.

**Figure 3.**
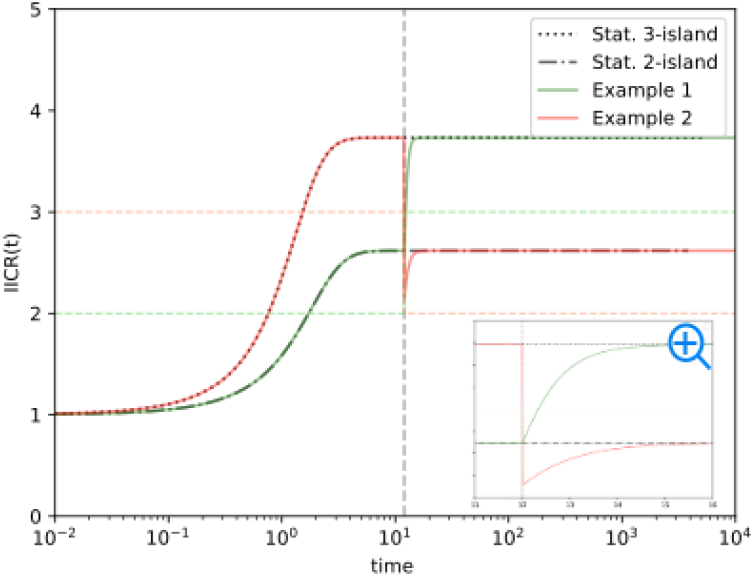
IICRs for the possible transitions between two and three islands. This figure represents the transitions of the reference example of Figure 2. The curves labeled “Stat.” are the stationary IICRs of the 2-island and 3-island models. All the migration rates between the islands are set to *M*_*ij*_ = 1. Time is measured in units of 2 × *N*_0_ × *t*_*gen*_ with *N*_0_ being the initial size of the population and *t*_*gen*_ the generation time. The horizontal dotted lines correspond to the evolution of the real population sizes for both backward deme addition (green) and backward deme suppression (red). The vertical dotted line indicates the time of the change in configuration (T = 12). The window provides a zoom-in view of the configuration change with time in natural scale.

This plot reveals that both non-stationary IICRs exhibit a consistent two-phase trajectory. During the initial phase, spanning from the present to the time of change, the curve aligns with the IICR of the respective stationary model: the 2-island model for backward deme addition and the 3-island model for backward deme suppression. Upon encountering the time of change, the IICR transitions abruptly from the initial stationary model to the other, thereafter following its trajectory for the remainder of the second phase.

However, there are distinct differences between the non-stationary IICRs at the time of the jump. In the case of backward deme addition, the curve rises continuously until it joins the plateau of the 3-island model. For backward deme suppression, the IICR decreases discontinuously below the plateau of the 2-island model before increasing backward in time and reaching its final plateau value. Even if the shape of the IICR is different in the case of the sampling in different islands (Supplementary Figure 9), we also retrieve the same two-phase trajectories that we just described.

From a forward perspective, the IICR thus appears to decrease in both cases, with this decrease seemingly “anticipating” the demographic change by starting before the actual change in configuration (proportionally to 4 × 2 × *N*_0_ × *t*_*gen*_). Furthermore, in the case of deme foundation forward in time, we observe a decrease in the IICR before it increases even though the population size is only increasing. Additionally, the size increase observed in the IICR does not appear to match the size of the new deme.

### Influence of the timing of a single demographic change: forward-time deme foundation

In this section, we analyze the impact of the timing of the change on IICRs. Employing the reference example depicted in Figure 2, we calculate the IICRs for potential transitions and systematically vary the timing of these changes, ranging from recent to ancient times.

Figure 4a illustrates the IICRs for the Example 2, with both genes sampled in the same island. (For the scenario where genes are sampled from different islands, refer to the Supplementary Figure 10.) Different types of IICRs emerge depending on the timing of the change. When the change occurs after the plateau of the second configuration has been reached backward in time (*i*.*e*. for large *T* values: *T* = 4, *T* = 12, *T* = 25), we observe similar curves to those described in the previous section, since we used a large *T* for the time of change. We observe for all these curves (*T* ≥ 4) a drop at the same IICR level below the plateau.

**Figure 4.**
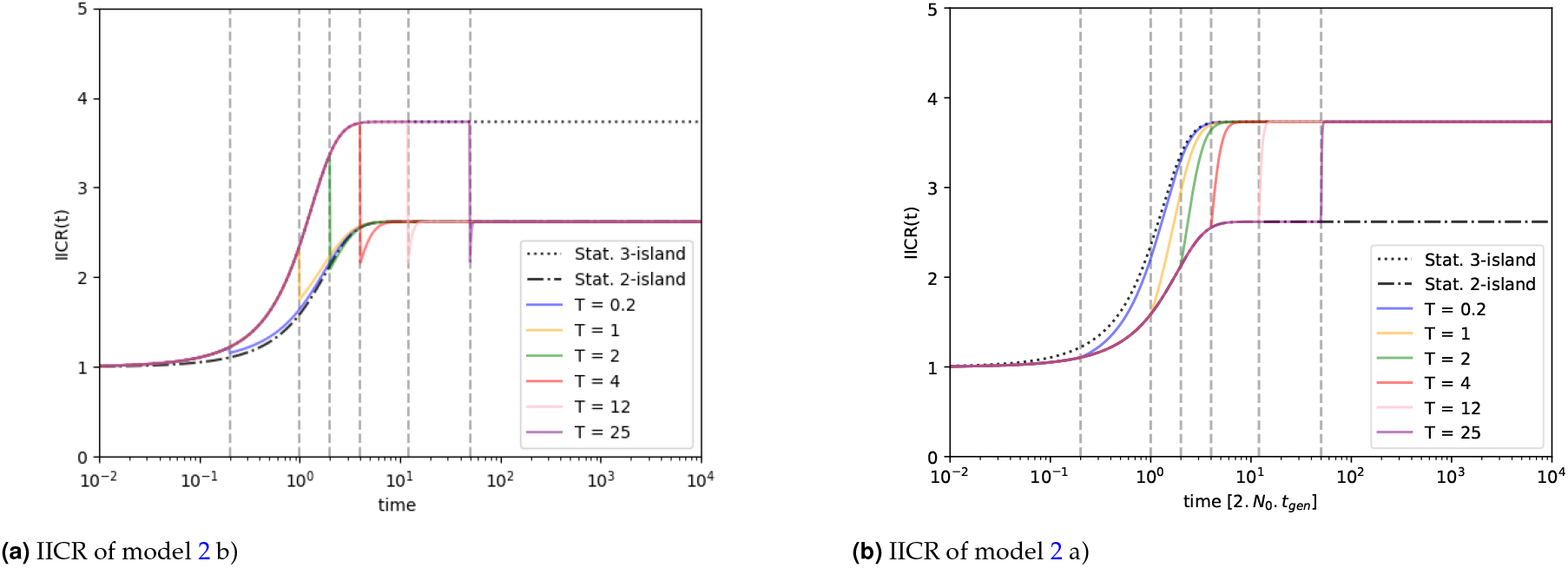
IICRs of both models of Example 2 (Figure 2) for different times of change in configuration (indicated by vertical dotted lines). The two genes are sampled in the same island. Time is measured in units of 2 × *N*_0_ × *t*_*gen*_ with *N*_0_ being the initial size of the population and *t*_*gen*_ the generation time. All the migration rates between the islands are set to *M*_*ij*_ = 1. The curves labeled “Stat.” are the stationary IICRs of the 2-island and 3-island models.

However, for more recent changes, particularly during the ascending phase towards the plateau (*T* = 1, *T* = 2), the step-wise decrease stops at a point located above the IICR of the second configuration before reaching it smoothly. The change becomes nearly undetectable when it occurs in the very recent past (e.g., *T* = 0.2), the non-stationary IICR seems to follow the stationary 3-island curve.

We note that if there is a deme extinction forward in time (Figure 4b), we will observe a clear step-wise decrease when the demographic event is older than the beginning of the plateau of the two-island model. If the change happens in a more recent time, the IICR will decrease forward in time and move from the 3-island to the 2-island IICR.

### Influence of the sampling scheme

Another factor influencing the IICR in the scenarios where a deme has been founded forward in time is whether we sample the recently founded demes or the other demes. As the founded deme does not exist in the past, we can suppose that the IICR of its lineages will differ from the other ones. Figure 5 displays the IICRs for two genes sampled in the same island in the case where the genes come either from a suppressed or non suppressed deme, illustrating both recent and ancient changes. (Refer to Supplementary Figure 11 for sampling in different islands.) We define ‘recent’ as a change occurring during the backward ascending phase of the IICR (or backward descending phase in the case of sampling from different islands), while ‘ancient’ refers to a change occurring when the plateau has been reached.

**Figure 5.**
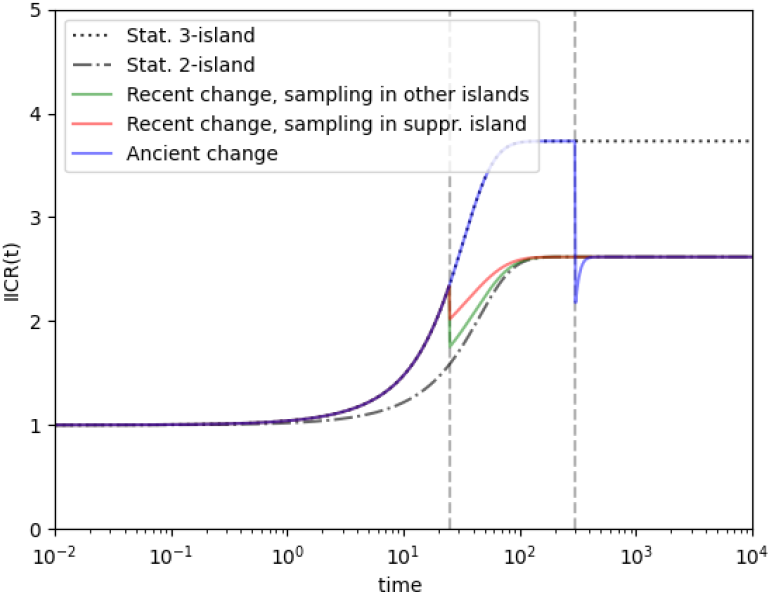
IICRs of Example 2 (model 2 b) for sampling in recently founded and ancient demes. In all cases, the two lineages are sampled in the same island. Time is measured in units of 2 × *N*_0_ × *t*_*gen*_ with *N*_0_ being the initial size of the population. All the migration rates between the islands are set to *M*_*ij*_ = 1. In one case the samples come from the deme that was founded and in the other case the genes come from one of the other two demes. The curves labeled ‘Stat.’ represent stationary IICRs of the 2-island and 3-island models. Vertical dotted lines indicate different times of configuration change.

As discussed in Section Influence of the timing of a single demographic change, if the change in structure occurs extremely close to the present (e.g., *T* = 0.2), the non-stationary IICR confounds with the stationary 3-island curve, making it difficult to observe any effect of the sampling. In the case of recent changes, we observe two distinct IICRs depending on the sample’s location. Specifically, when both lineages originate from the disappearing island (indicated by the red curve on the graph), the drop in the IICR is less pronounced compared to other scenarios (indicated by the green curve).

After the plateau has been reached, the effect of sampling becomes indiscernible, and all the curves are confounded into the blue curve. Similar observations emerge when examining the curves for sampling in different islands (refer to Supplementary Figure 11).

From a forward perspective, we observe that if the change is recent enough, it influences the “bottleneck” observed just before the change in configuration. When we sample in the newly founded deme, the bottleneck is less pronounced than when we sample in any other deme. We thus observe two rather non intuitive results: if a deme has always been present, the IICR suggests a stronger decrease (forward in time) if we sample in that deme rather than in a deme that has recently been founded, and these changes start before the foundation of the deme in a forward perspective.

### Influence of the migration rate

We varied the migration rates from *M*_*ij*_ = 0.1 to *M*_*ij*_ = 100 in the migration matrix. The IICRs for the transition from a 3- to 2-island models (backward) with the sample taken in the same island (island 1) are depicted in Figure 6. (Refer to the Supplementary Figure 12 for the scenario involving sampling in different islands.)

**Figure 6.**
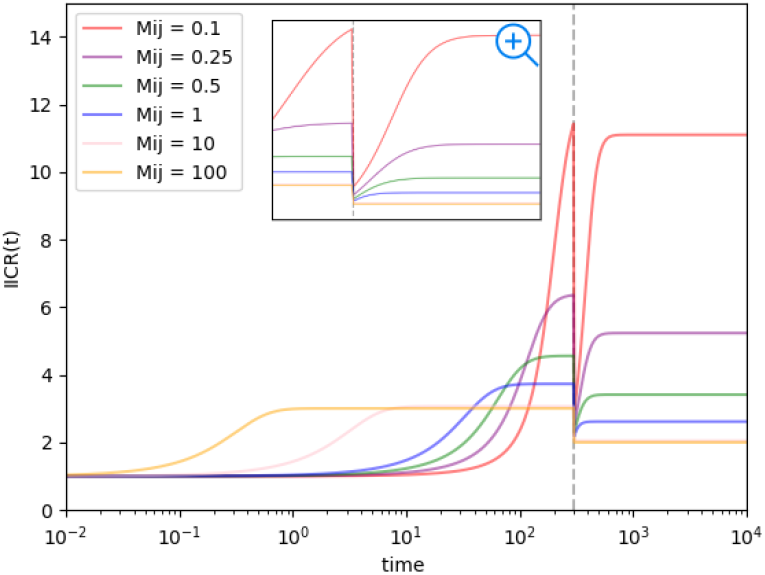
IICRs of Example 2 (model 2 b) for different migration rates. Both lineages are sampled from the same island (island 1, non-founded). The vertical dotted line indicates the time of the configuration change (T = 12). Time is measured in units of 2 × *N*_0_ × *t*_*gen*_ with *N*_0_ being the initial size of the population and *t*_*gen*_ the generation time. The window provides a zoom-in view of the configuration change and uses a natural time scale.

Lower migration rates result in higher plateaus occurring in more ancient times, consistent with previous observations (e.g., Mazet *et al*. (2016)). These lower migration rates lead to significant drops below the plateau of the 2-island model, deeper than those induced by higher migration rates. For high migration rates, the jump resembles a step function. The drop indicating a change in configuration becomes increasingly indistinguishable as the plateaus of the two models get closer to each other. The zoom-in view shows that the curves do not drop to the same point. We still observe that the curve starts to decrease several generations before the change in configuration except in the case of high migration rates (*i*.*e. M*_*ij*_ = 100).

### Extension to more complex examples

We can now examine the IICRs of more complex processes that may be seen as simple “real-life” biological processes, possibly shedding light on patterns observed in published PSMC curves derived from real data. In Figure 7, we present two examples of IICRs, one depicting a step-wise colonization process and the other illustrating a split process.

**Figure 7.**
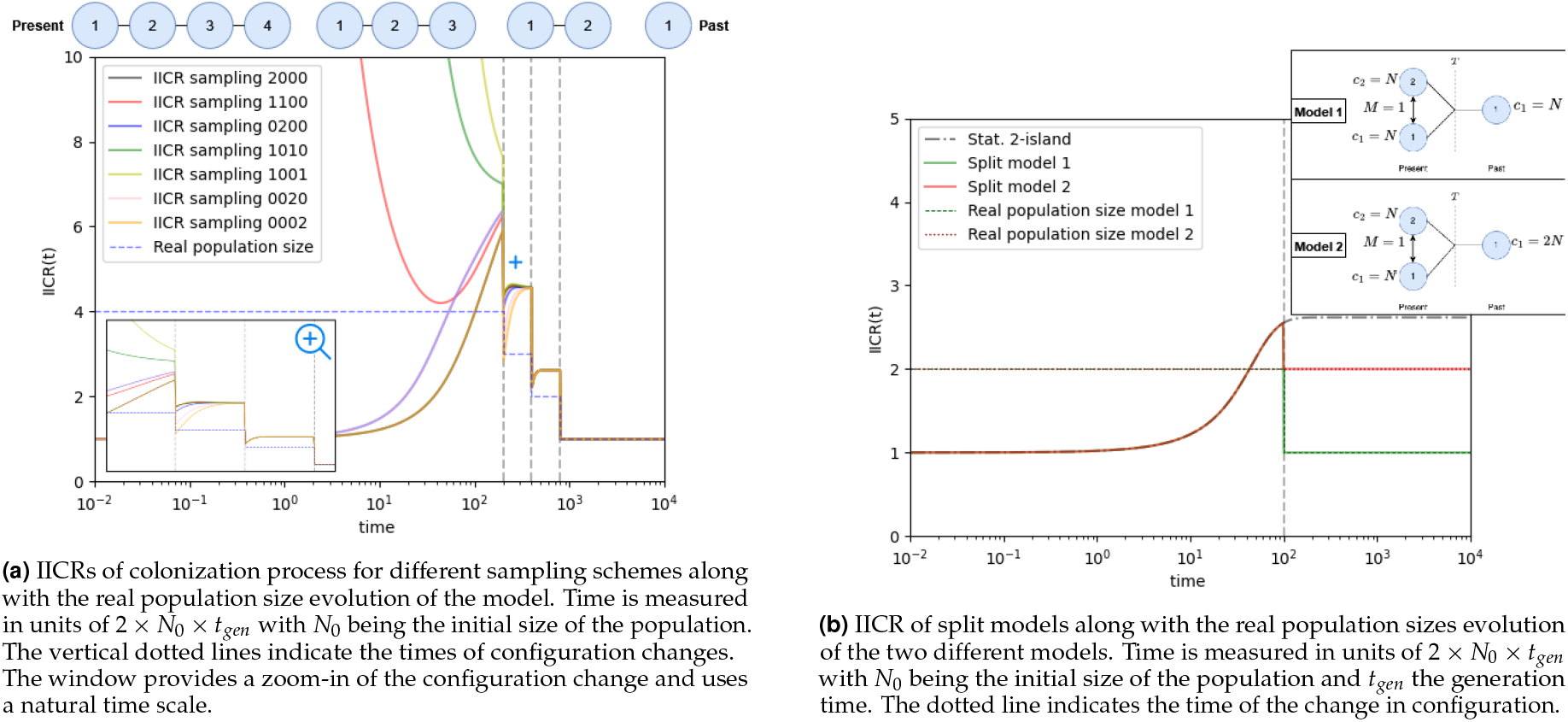
IICRs of more complex examples with representations of their corresponding structured models.

**Figure 8.**
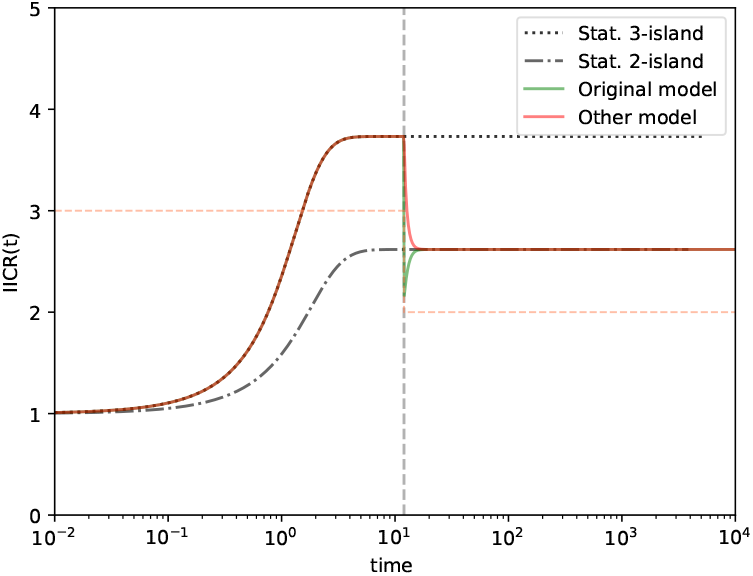
IICR of the Example 2 of the reference example of Figure 2. The curves labeled “Stat.” are stationary IICRs of 2-island and 3-island model. Time is measured in units of 2 × *N*_0_ × *t*_*gen*_ with *N*_0_ being the initial size of the population and *t*_*gen*_ the generation time. The dotted line corresponds to the evolution of the real population size. The vertical dotted line indicates the time of the change in configuration.

#### Colonization process

Theoretically, our model allows for an unlimited number of configuration changes that are achieved by constructing a corresponding glue matrix for each transition. Here, we considered a stepwise linear colonization model, where a single panmictic population colonizes a new deme to which it is connected by gene flow and from which a new deme is founded. The process is repeated until we have four populations connected as a linear stepping-stone model. We computed the IICR for several sampling schemes to illustrate the diversity of IICR curves that can be obtained (see Figure 7a). We note that the IICR exhibits a regular increase from the past to the present as the number of islands increases but this increase is also characterized by the noticeable drops beneath the plateaus that were observed in Example 2. Interestingly, these drops are stronger in more recent times. We also note that the IICR exhibits plateaus that are distinct from the actual size of the population. We can also see the influence of the sampling, with the IICR curves changing with the age of the sampled island. Altogether, the IICR exhibits a general increase in the IICR, forward in time, with the increase in the number of demes, and a final decrease typical of an *n*-island stationary IICR.

#### Split/fragmentation process

We studied a particular split model called isolation-with-migration model whereby a single panmictic population divides *T* generations ago into two demes of size *N* that are connected by gene flow. In one model, the population size of the ancestral population is *N* and in the other it is 2*N*. Two IICRs are depicted in Figure 7b, for a case where the split time takes place when the plateau has been reached (backwards) by the stationary IICR for the two-island model. We observe that the IICRs exhibit simple step functions, without any decrease below the plateau that we observed in the previous models. The plateau’s value corresponds to the size of the ancestral population, as expected since the ancestral population is panmictic.

## Discussion

The IICR was introduced by Mazet *et al*. (2016) in an attempt to understand the properties of genetic data under structured models. This work followed several studies that showed that spurious population size changes can be detected, dated and quantified when populations are structured and when structure is assumed to be negligible during the inferential process (Wakeley 1999; Beaumont 2004; Chikhi *et al*. 2010). The IICR was crucial in showing quantitatively how a stationary *n*-island model would generate a specific S-shaped population size change from an ancient stationary population to a recent stationary population whose size would be the size of the local sampled deme. The IICR was originally defined for two genes and analytical results were obtained for the *n*-island model. For more complex model it was shown that IICR could be computed using coalescent simulators such as ms (Hudson 2002) or msprime (Kelleher *et al*. 2016; Chikhi *et al*. 2018). Rodríguez *et al*. (2018) showed that a general matrix approach could be used to compute the IICR for non stationary models for which the total number of states did not change. Here we extended their work by showing that the computation of IICRs for non-stationary models with changes in the number of states is possible thanks to the introduction of glue matrices. We also proposed a general method for computing these glue matrices based on relatively simple hypotheses: the genes of a newly-founded island (in a forward perspective) come from all the islands that are connected to this new deme, and the genes of an extinct island cannot be sampled in the present and do not contribute to its genealogy. Finally, we analyzed the resulting IICR curves.

As in previous studies, we have shown that under the dynamical structured models examined here, the IICR curves can significantly differ from the actual population size. In the recent past, assuming constant-size models and sampling in the same island, the IICR tends to reflect the deme size but then significantly diverges from the total population size as we go back in time. In some cases, the IICR may even exhibit trends that are contrary to the actual population size changes. Therefore, our results confirm that the PSMC and MSMC methods, which infer the IICR, should not be interpreted as direct indicators of population size changes unless additional evidence supports such interpretations.

Our modeling approach produced IICR curves with qualitatively intuitive patterns, but also with several non-intuitive patterns even considering our current understanding of IICRs. A particularly non-intuitive notable case is that of deme suppression backward in time, where we observed two surprising features. First, the IICR curve descended below the expected plateau before reaching it. Second, from a forward point of view, the IICR curve drops below the lower plateau before any demographic event takes place, then rises towards the expected IICR curve, which subsequently decreases due to population structure. This indicates that the IICR curve is highly disconnected from the actual dynamics of the total population size. Additionally, the vertical drop below the plateau observed is a new result not seen in previous studies in the context of a neutral model. This phenomenon can be explained by our modeling choice, as expressed in the glue matrices. In our model, genes located in islands that do not disappear stay where they are, while genes from a disappearing deme are equally distributed among the neighboring islands that continue to exist. Consequently, genes that were in different islands (impossible coalescence states) can end up in the same island (possible coalescence state). This effectively increases the probability of coalescence instantaneously at the time of the change, resulting in a drop in the inverse probability of coalescence, and thus the IICR. This feature may not necessarily be present with a different modeling approach, as demonstrated in Section Supplementary Material.

Our findings also align with intuitive expectations in certain scenarios. Specifically, when new islands are founded forward in time (not too recently), the IICR increases step by step, reflecting the overall increase in total population size. Conversely, when demes go extinct (not too recently), the IICR decreases, along with the total population size. Even if the increase (or decrease) is not proportional to the size of the deme that is added/suppressed, this intuitive result reinforces the idea that the IICR can capture some of the effects of population expansion or contraction. This highlights the utility of IICR as a tool for understanding population growth and decline, provided that the underlying population structure is accurately accounted for.

An important aspect to note is the versatility of glue matrices, which can be computed according to different modeling hypotheses. We show in Supplementary Material that the drop below the plateau can be eliminated by adopting alternative hypotheses, considering genes by pair at the time of change. In this scenario, lineages in the same island can only move to states in which they remains in the same island, while lineages in different islands continue to stay in separate islands. However, we could not find a clear biological significance for these hypotheses. This matrix primarily serves as proof that changing the value of *n* in the simplified *n*-island matrix is not an appropriate modeling approach. It is also conceivable that some glue matrices produce a hump before reaching the plateau.

Observing different drops induced by various sampling strategies (Figures 5, 11), especially when one sample includes a lineage from an extinct island while another does not, could potentially aid in identifying a recent change in configuration. However, this is only feasible if the change is not too recent or too ancient, as the two curves would be confounded.

Furthermore, an accurate method for estimating the IICR based on real data is necessary, as the difference between the two curves may not be significant enough.

However, some of these artifacts may not be visible on PSMC curves because they occur on a too small a timescale compared to the time intervals used in the time-discretization of PSMC (see Supplementary Material). Further work is needed to address this, involving the simulation of genomes and the application of the PSMC method to data generated under the scenarios discussed here.

Expanding the non-stationary structured coalescent framework offers a novel tool for inferring more complex structured models. This extension allows for the consideration of factors such as changes in the number of demes. By computing IICRs for non-stationary models, we could design a new inferential tool similar to SNIF (Arredondo *et al*. 2021) that would allow changes in the number of demes. Such a tool would be especially useful for studying split models and inferring key events, such as the timing of population splits or colonization processes. However, the parameter space becomes significantly larger, necessitating a focused effort to precisely define the inference problem. This can be achieved by applying appropriate constraints such as limiting the number of configuration changes, or introducing simplifying assumptions about the population structure, like assuming a stepping-stone model, a continent-island model, or other structured models commonly used in population genetics. Stepping-stone models are particularly interesting to capture the spatial structure of our population.

Going even further, we could use multiple IICRs to infer parameters of our models. Indeed, we can easily extend the glue matrices to compute IICRs of samples of size *k* by just defining a new mapping of states that takes into account all the possible states arising from the larger sample size.

## Acknowledgments

We would like to thank the collaborators from the DevOCGen project for the comments that helped improve the work presented in this manuscript. We thank also Willy Rodriguez for the Python code that allows the computation of IICR using Markov chains and that was extended in this work.

## Data availability

The code to compute non-stationary IICRs and reproduce the figures of the article is available on Github at https://github.com/aljounia/Theoretical_IICR_with_glue_matrices_generator.

## Appendix: Supplementary Material

### What is the probabilistic meaning of glue matrices?

Let *α*(*t*) denote the time-homogeneous continuous time Markov chain modeling the structured coalescent of our sample of two lineages. It gives the ancestral lineages configuration at time *t* in the past, so for example *α*(0) denotes the present sampling configuration. It is a right-continuous process, meaning in particular that *α*(*T*) is the configuration at time *T* once the change in model parameters has happened, and so *α*(*T*)≠ *α*(*T*^−^).

If we have two possible configurations of the lineages (*ij*) and (*kl*), we know that the element at the index (*n*_(*ij*)_, *n*_(*kl*)_) of a certain transition matrix *P*_*t*_ give us the following conditional probability ℙ [*α*(*t*) = (*kl*) | *α*(0) = (*ij*)], ∀*t* ≥ 0 (*n*_(*ij*)_ is the index of the row of configuration (*ij*) and *n*_(*kl*)_ the index of the column of configuration (*kl*)).

To better understand the construction of 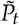, we will explicit the product 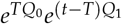 in terms of conditional probabilities. First, we note that 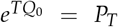 and 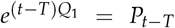. The element (*n*_(*ij*)_, *n*_(*kl*)_) of the matrix *P*_*T*_ give us the probability ℙ [*α*(*T*) = (*kl*) | *α*(0) = (*ij*)]. The element (*n*_(*ij*)_, *n*_(*kl*)_) of the matrix *P*_*t*−*T*_ give us the probability ℙ [*α*(*t* − *T*) = (*kl*) | *α*(0) = (*ij*)] which is equivalent to ℙ [*α*(*t*) = (*kl*) | *α*(*T*) = (*ij*)] by the property of time-homogeneity of the Markov chain *α*(*t*).

In the case of a parameter change without changing the number of demes, we have then:

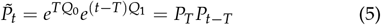

which can be translated as:

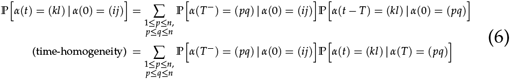

If we change the number of demes, we must use the glue matrix *G* to allow the multiplication of the exponentials. We construct this glue matrix such as the element at index (*n*_(*ij*)_, *n*_(*kl*)_) corresponds to the instantaneous probability ℙ[(*ij*) → (*kl*)] at the time of the transition *T*. More explicitly, at time *T*^−^ if the lineages are in the state (*ij*), they will have a probability

ℙ [*α*(*T*) = (*kl*) | *α*(*T*^−^) = (*ij*)] to be in the state (*kl*) at time *T*.

We have then:

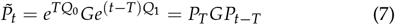

which gives:

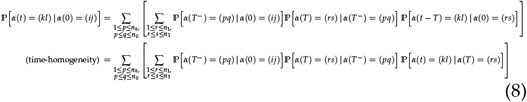

### Other possible modeling

In this alternative modeling, we only match states where the two lineages are in the same deme with the ones where the two lineages are in the same deme. Different-deme states match different-deme states. When the two lineages are in the newly created island, we assume that they can come from all the “same-deme” states with the same probability. It can be translated as:

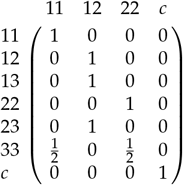

The IICR curve resulting is shown in Figure 8.

If we use the simplified transition matrices of *n*-island models (described in section The structured coalescent and the IICR) and simply change the *n* to modify the number of demes, we do not need a glue matrix and we can directly compute the IICR. In this case, we find the same IICR as for the modelling we described in this section.

## Appendix: Results Supplementary Figures

**Figure 9.**
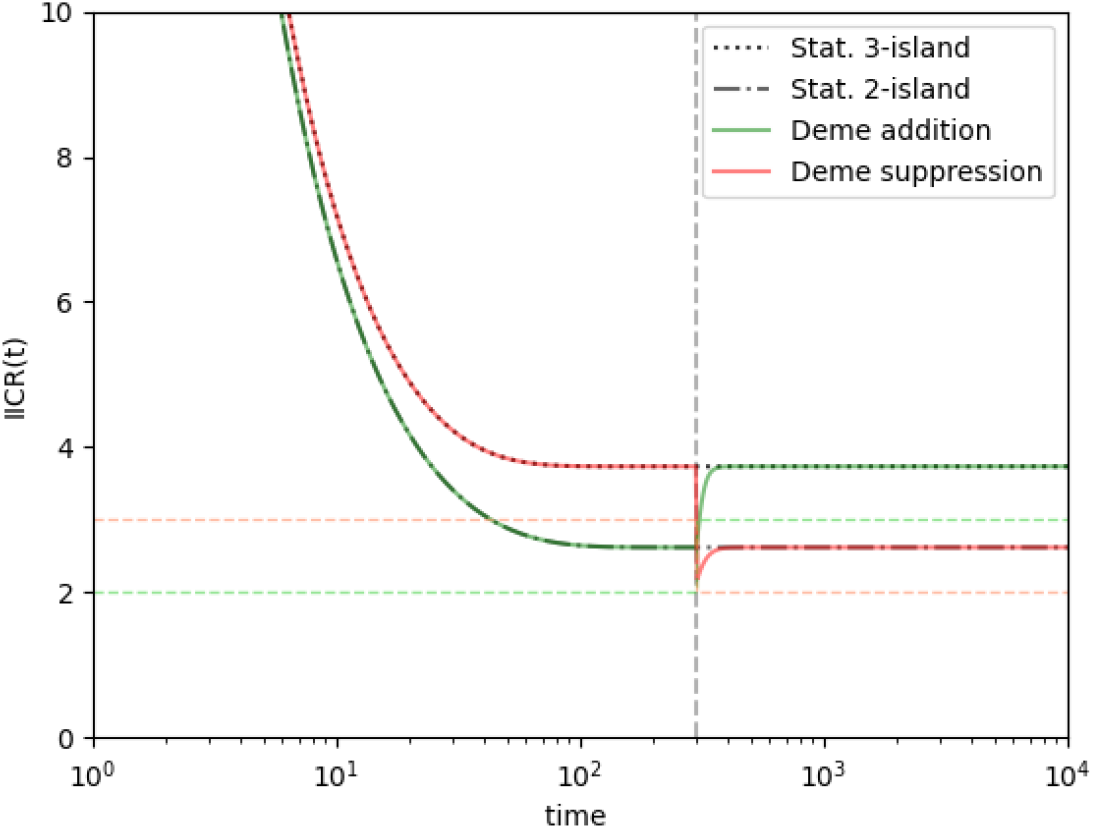
IICRs of the two possible transitions of the reference example of Figure 2. The curves labeled “Stat.” are stationary IICRs of 2-island and 3-island model. Time is measured in units of 2 × *N*_0_ × *t*_*gen*_ with *N*_0_ being the initial size of the population and *t*_*gen*_ the generation time. All the migration rates are set to *M*_*ij*_ = 1. The dotted lines correspond to the evolution of the real population sizes for both deme addition (green) and deme suppression (red). The vertical dotted line indicates the time of the change in configuration.

**Figure 10.**
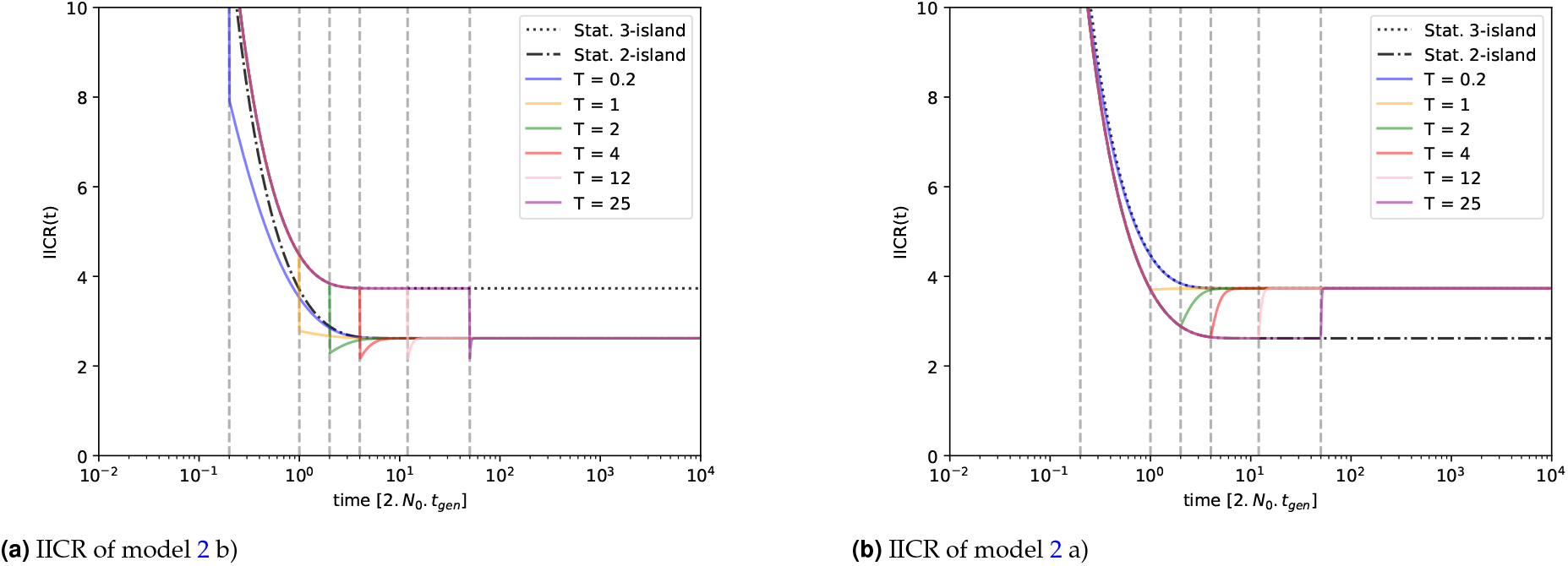
IICRs of both models of the Example 2 (2) for different times of model change. The two genes are sampled in different islands. Time is measured in units of 2 × *N*_0_ × *t*_*gen*_ with *N*_0_ being the initial size of the population and *t*_*gen*_ the generation time. All the migration rates between the islands are set to *M*_*ij*_ = 1. The curves labeled “Stat.” are the stationary IICRs of the 2-island and 3-island models. The vertical dotted lines indicate the different times of change in configuration.

**Figure 11.**
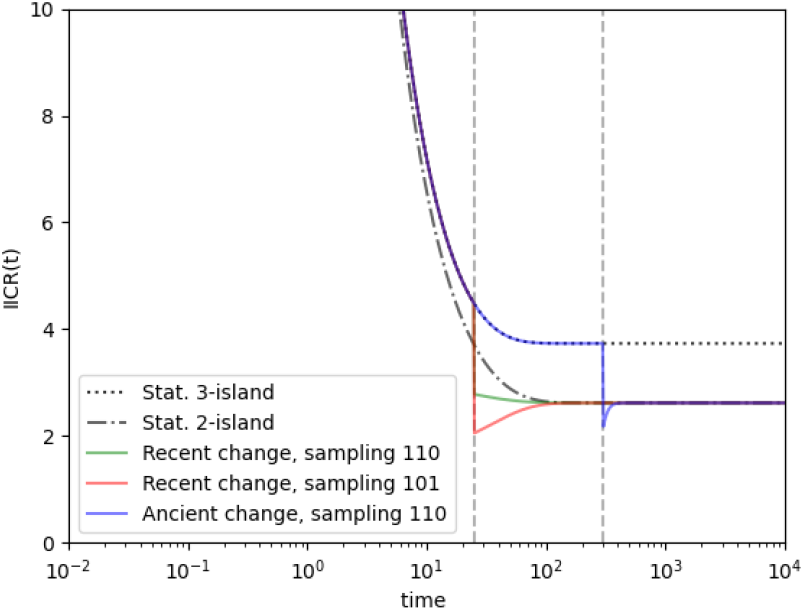
IICRs of the transition from 3- to 2-island model in a backward point of view (model 2 b) for different sampling schemes. Lineages are sampled in different islands. The curves labeled ‘Stat.’ represent stationary IICRs of the 2-island and 3-island models. Time is measured in units of 2 × *N*_0_ × *t*_*gen*_ with *N*_0_ being the initial size of the population and *t*_*gen*_ the generation time. All the migration rates are set to *M*_*ij*_ = 1. Vertical dotted lines indicate different times of configuration change.

**Figure 12.**
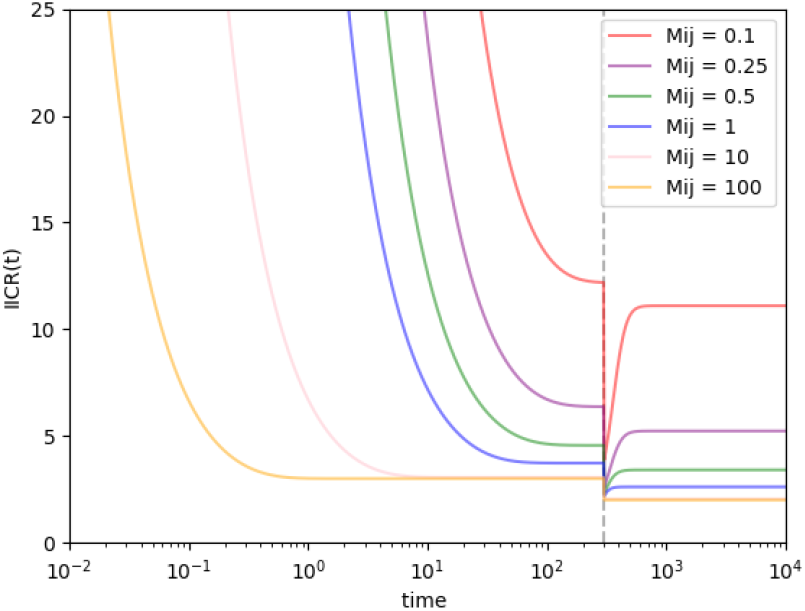
IICRs of the transition from 3- to 2-island model in a backward point of view (model 2 b) for different migration rates. Lineages are sampled in different islands. Time is measured in units of 2 × *N*_0_ × *t*_*gen*_ with *N*_0_ being the initial size of the population and *t*_*gen*_ the generation time. All the migration rates are set to *M*_*ij*_ = 1. The vertical dotted line indicates the time of the configuration change. The window provides a zoom-in view of the configuration change.

